# A 128-ch area-efficient neurochemical-sensing front-end for FSCV recordings of dopamine

**DOI:** 10.1101/2023.06.20.545834

**Authors:** Kevin A. White, Mahdieh Darroudi, Jinwoo Park, Brian N. Kim

## Abstract

Neurochemical recordings rely on electrochemical reactions of electroactive neurotransmitters such as dopamine, serotonin, and norepinephrine. This electrochemical technique allows for highly sensitive monitoring of neurotransmitters in the brain. Traditionally, single-channel carbon-fiber microelectrode recordings have been considered the gold standard method. However, an alternative approach involves the use of a microelectrode array, which enables high spatiotemporal resolution imaging of electroactive neurotransmitters. To enable neurochemical imaging using a microelectrode array, the development of a high-density current-sensing microchip is necessary. Here, a neurochemical microchip is introduced, featuring a 128-channel current sensing front-end capable of supporting 128 parallel neurochemical measurements. The designed amplifier array employs a highly scalable resistive feedback transimpedance amplifier design. This design allows for a large neurochemical dynamic range of ±5µA with a noise performance as low as 0.22nA_RMS_. With the integration of this microchip, *in vivo* neurochemical imaging of dopamine can be achieved with high spatiotemporal resolution.

## I. Introduction

High spatiotemporal imaging of dopamine in the brain is essential to study its complex interactions and functions in different regions and neural circuits. Dopamine (DA) is a key neurotransmitter that plays a crucial role in a wide range of neurological processes, including motivation, reward, and addiction. By imaging dopamine in real-time and with high spatial resolution, we can help elucidate the neurological function of neurochemicals in the central nervous system [1-5]. Moreover, high spatiotemporal imaging of dopamine is critical for investigating dopamine-related disorders such as Parkinson’s disease, schizophrenia, and Alzheimer’s disease. Precise localization of dopamine release and uptake dynamics could aid in the identification of potential biomarkers or therapeutic targets for these disorders.

Electrochemical recordings of dopamine are widely used in neuroscience research as they provide real-time, highly sensitive measurements of dopamine release and uptake dynamics in the brain. These recordings are based on the electrochemical reactions of dopamine and other electroactive neurotransmitters at the surface of an electrode, which generates a current proportional to the amount of neurotransmitter present. Single-channel carbon-fiber microelectrode (CFE) recordings have been the gold standard for neurochemical recordings, but they are limited in their ability to capture spatial variations in neurotransmitter release. Current-sensing analog front ends are used as bioinstrumentation to apply a potential to an electrode to induce redox reactions while measuring the redox current (Fig. 1). Although a single carbon-fiber microelectrode is commonly utilized for in vivo monitoring of dopamine activity through fast-scan cyclic voltammetry (FSCV) (Fig. 1b) [1, 6-11], a microelectrode array can offer a high spatiotemporal resolution by gathering neurochemical activity from a particular brain region to generate neurotransmitter mapping. To enable the operation and recording of a microelectrode array with a large number of electrodes (hundreds), a microchip containing an equal number of amplifiers is necessary to facilitate highly parallel electrochemical recordings [12]. With the development of high-density microchips for processing and amplifying the small redox signals generated by these microelectrodes, electrochemical recordings of dopamine can provide a powerful tool for investigating the complex dynamics of dopamine signaling in the brain and its role in neurological diseases.

**Fig. 1.**
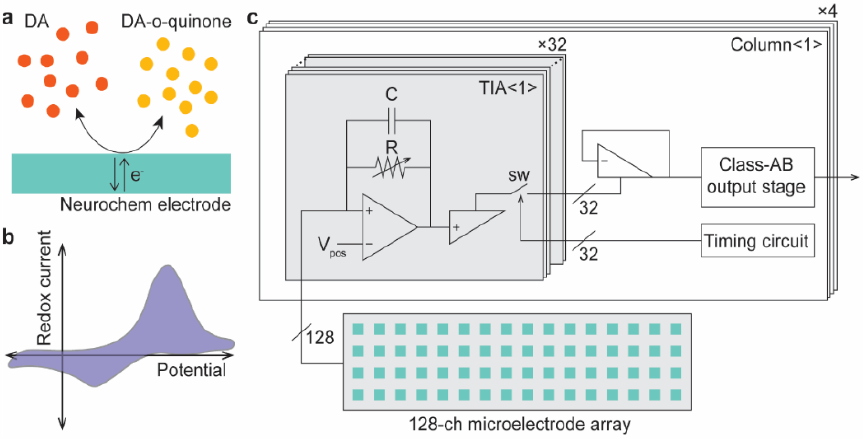
Dopamine (DA) detection using 128-ch neurochemical front-end and fast-scan cyclic voltammetry (FSCV). (a) Dopamine is detected by redox reactions at an electrode using FSCV, wherein molecules lose electrons (oxidation) or gain electrons (reduction). (b) FSCV recordings provide identification of a molecule, as well as quantity of molecules, by the location and intensity of redox peaks vs. the electrode potential. (c) The neurochemical sensing system incorporates a 128-ch neurochemical front-end for FSCV recordings using resistive-feedback transimpedance amplifiers.

Typically, resistive-feedback or integrating amplifier topologies are used to design analog front ends for electrochemical recordings. An integrating amplifier uses an operational amplifier (OPA) to regulate the electrode’s potential and collects the electrode’s current through a capacitor. The voltage change across the capacitor is then periodically read to monitor the current detected by the electrode. The main advantage of integrating amplifiers is their low noise characteristics [12-19]; however, for large-dynamic range applications such as *in vivo* neurochemical recordings, the capacitor size can become too large, and integrating hundreds of amplifiers with a small and scalable silicon footprint can be challenging. For example, to achieve a ±5 µA dynamic range with a 20 kHz sampling rate, the integrating capacitor must be around 150 pF. A CMOS potentiostat using a 50-fF integrative capacitor was only able to record between ±1.5 nA [13]. Another CMOS potentiostat using 50 to 150 fF integrative capacitor also has a limited dynamic range of 60 nA [20]. Although it is possible to integrate a few 150 pF capacitors using CMOS technology, it is challenging to integrate 100s of amplifiers. Current division or current subtraction can be used to decrease the required capacitance [21-23]; however, they come with a cost of power, area, or noise performance. An analog front-end with analog background subtraction was able to achieve high bandwidth (5 kHz) and low noise level (26.5 pA_RMS_), however, the analog front-end size was relatively larger (0.256 mm^2^) compared to other works that can achieve similar dynamic range [23]. Another subtraction-based circuit had a smaller footprint (0.06 mm^2^), however, it suffered from a significant noise floor (10 pA/√Hz). Resistive feedback-based transimpedance amplifiers (TIA) can be suitable for large-dynamic range applications because the feedback resistors (around 1 MΩ and smaller) can be relatively smaller compared to equivalent capacitors (around 200 µm × 30 µm for 1 MΩ in a standard 0.35 µm process) [24].

In this paper, we present a high-density neurochemical-sensing analog front-end microchip that integrates 128 resistive TIAs with multiplexers to support fully parallel 128-ch FSCV recordings of electrochemical activity (Fig. 1c). The paper is organized as follows. In Section II, the design of the neurochemical-sensing microchip and the circuit implementation are described. The characterization of gain, noise, and bandwidth performances are discussed in Section III. Section IV describes the carbon-fiber electrode fabrication and dopamine recordings using the neurochemical microchip. Finally, the conclusions and discussions are described in Section V.

## II. Highly scalable neurochemical-sensing amplifiers

To design a large array of TIAs, it is critical to develop a TIA that occupies a small footprint. While both resisitive and capacitive feedbacks are commonly used, we consider both options to minmize the design area.

### A. Highly scalable transimpedance amplifier design

In a standard 0.35 µm process, a poly capacitor has a capacitance density of ∼0.90 fF/µm^2^. Instead, an n-well resistor is ∼1,000 Ω per square and poly resistor is 50 per square. Given that the minimum width of a n-well resistors is 3 µm, the resistance density can be calculated to be 110 – 120 Ω/µm^2^. This translates to a gain density of 110 – 120 V/A/µm^2^ (Fig. 2a). Therefore, using resistive feedback, the gain increases linearly with the invested area.

**Fig. 2.**
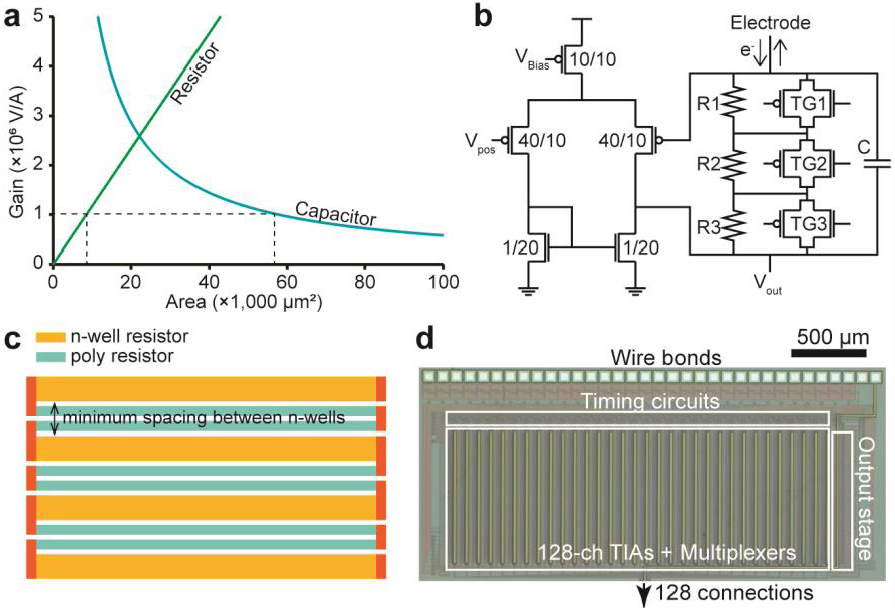
Resistive feedback TIAs for 128-ch neurochemical sensing using FSCV. (a) Optimal transimpedance gain is in the range of range of 0.1 – 1 µA/V. To achieve this, a feedback resistor-based TIA can provide comparable gain to an integration capacitor-based TIA with lower area investment in layout. (b) The embedded transmission gate switches TG1 – TG3 enable configurable TIA gain in tandem with the segmented feedback resistance. The feedback resistance directly determines the gain of the TIA and gain on the scale of 100 kΩ – 1 MΩ can be configured. (c) To achieve maximum area efficiency when creating the layout, poly resistors are used in the spacing required between the n-well resistors for maximum resistance density. (d) The current sensing front-end has 128 parallel channels for neurochemical detection with the presented feedback resistor TIA arranged in a 4 × 32 array.

In an integrating amplifier that uses capacitive feedback, the transimpedance gain is set by the integration time and capacitance:

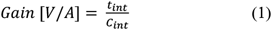

where *C*_*int*_ is the value of the integration capacitor, *t*_*int*_ is the integration duration. Assuming a sampling rate of 20 kS/s, the highest integration time that can be allocated is 50 µs. The gain determined by poly capacitor is ∼6×10^10^ V·µm^2^/A. When the targeted transimpedance gain is 1×10^6^ V/A, the resistive feedback requires substantially less space (∼9,000 µm^2^) compared to that of capacitive feedback (∼60,000 µm^2^) (Fig. 2a). Based on the gain-area plot, the integration capacitor becomes more area efficient than the resistor when the desired gain is beyond 2.6 ×10^6^ V/A. For FSCV measurements coupled with microelectrodes, the targeted transimpedance gain is typically between 0.1 – 1 µA/V, corresponding to 100 kΩ – 1 MΩ. Therefore, the resistive TIA is significantly more area efficient.

Our TIA design is based on resistive feedback in which the incoming redox current is multiplied by the resistance and directly changes the amplifier’s output voltage (Fig. 2b). The feedback resistance is divided into three resistors to enable programmable transimpedance gain. Each segmented resistor is linked to a parallel transmission gate (TG1 – 3), which allows for adjustable transimpedance gain. The three transmission gate switches can be configured to produce 2^3^ different gain settings. Because n-well resistors achieve highest resistance density compared to other types of resistors present in the CMOS technology, n-well resistors are the preferred option for the highly scalable TIA design. However, the design rule sets a minimum spacing (typically equivalent to the minimum width of a n-well resistor) between n-wells with different potentials which adds a large overhead in integrating multiple segments of n-well resistors in series. This rule is to avoid significant current flow between n-wells. To maximize the resistance density, poly resistors can be added within the spacings (Fig. 2c). Although the resistance per square for poly resistors is far smaller compared to that of n-well resistors, because the minimum width is smaller (below 1 µm^2^), the resistance density is ∼60 Ω/µm^2^, nearly half of that of n-well resistors. The three feedback resistors are made up of a combination of n-well resistors and poly resistors (as shown in Fig. 2c). This configuration provides the highest resistor density per area without violating layout design rules.

The transistor dimensions are selected to optimize the area efficiency while preserving gain, bandwidth, power efficiency, and uniformity based on Monte Carlo simulation (Fig. 2b). The feedback capacitor restricts the bandwidth to a range of 10 – 100 kHz. The oxidation/reduction current flows through the feedback resistor, creating the transimpedance gain at the OPA’s output. Each TIA occupies 220 µm × 80 µm (0.0176 mm^2^) of space. A total of 128 TIAs occupies only 2.25 mm^2^ silicon space (Fig. 2b).

### B. Time-division multiplexing

To reduce the number of outputs, the amplifiers’ outputs are combined in groups of 32 as columns using time-division multiplexing. The system comprises 4 columns with a total of 128 fully parallel TIAs. The time-division multiplexing is based on a half-shared amplifier that connects individual TIA’s outputs to the output stages using switches [12, 15, 17]. Each TIA contains an embedded non-inverting OPA (three transistors), which links to an inverting OPA in each column via a switch (Fig. 3a). When Row1 is low (active low), the other row switches (Row2 – Row32) are high so that only In1 is the unity gain input to the multiplexer’s output, MuxOut. In a 50-µs interval, all 32 inputs (In1 – In32) are multiplexed to MuxOut by staggering the switches (Row1 – Row32). The output in Fig. 3b shows the multiplexed output where a TIA connected to In1 (TIA #1) has an input current of -1µA or 2µA. For this test, the rest of TIAs have zero input current and therefore results in a flat output.

**Fig. 3.**
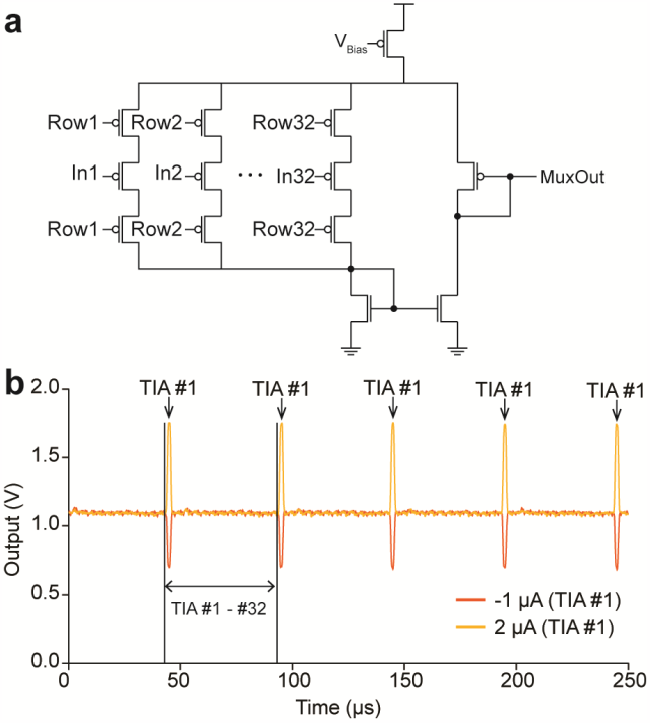
(a) The multiplexer is a half-shared structure. The non-inverting side is embedded in each amplifier and the inverting side is shared as a multiplexer output. (b) The multiplexer uses time-division multiplexing.

### C. System integration

The system consists of 128 TIAs, multiplexers, timing circuits, and output stages (Fig. 1c). As the outputs from this system is sampled using external analog-to-digital converters (ADC), class-AB output stages are designed within the system to drive large capacitive loads originating from the ADC’s input capacitance and the line capacitance. The timing circuit produces all the sub-clocks required to run this multiplexing scheme. The amplifier array is fabricated using a standard 2-poly 4-metal 0.35-µm process (Fig. 2). This process is chosen because of its wide voltage range. The amplifier array, timing circuit, output stage and wire bonds occupy 3.05 mm × 1.37 mm.

## III. Performance AND Characteristics

Transimpedance gain, bandwidth, noise, and crosstalk of the TIA is evaluated in this section. For all tests, USB-6363 (National Instruments) is used for analog data acquisition. Current signals are generated using either USB-6363 DAQ for alternating current or a source/measure unit (B2901A, Keysight) for constant current. For noise measurements, the chip is powered using battery-powered voltage regulators to minimize the contribution of line frequency. For all data acquisitions, a custom-designed LabVIEW program is used. In this program, the clock source that governs the data sampling is synchronized with that of digital-to-analog converters (DACs) so that the artificial signals being injected into the amplifier have a sharply defined frequency component. This is beneficial in analyzing the frequency response of the amplifiers.

### A. Measured transimpedance gain

To determine the gain of the current sensing front-end, an external current is introduced into the amplifier and the resulting change in output voltage is recorded (Fig. 4a). Across the dynamic range shown, the amplifier exhibits a highly linear response in four gain settings, with an R^2^ value of 0.999 for all gain settings. The dynamic range can vary from as high as ± 5 μA (Gain 4) to as low as ± 650 nA (Gain 1). Furthermore, the uniformity of the transimpedance gain across the array is evaluated. Any mismatch in the dimensions of devices in each TIA can contribute to non-uniformity in performance characteristics. The array demonstrates a consistent gain across the 128 amplifiers in the array (as Fig. 4b), with measured gains of 1.16 ± 0.04 MΩ, 618.93 ± 13.3 kΩ, 314 ± 9.63 kΩ, 127 ± 2.17 kΩ (mean ± SD) for Gain 1, Gain 2, Gain 3, and Gain 4, respectively. The transimpedance gains plotted in histograms show guassian distributions for each gain setting (Fig. 4b). Because this deviation is systematic, gains of each amplifier can be measured during a calibration step, and subsequantly applied individually to each amplifier’s recordings to accurately represent the measured current.

**Fig. 4.**
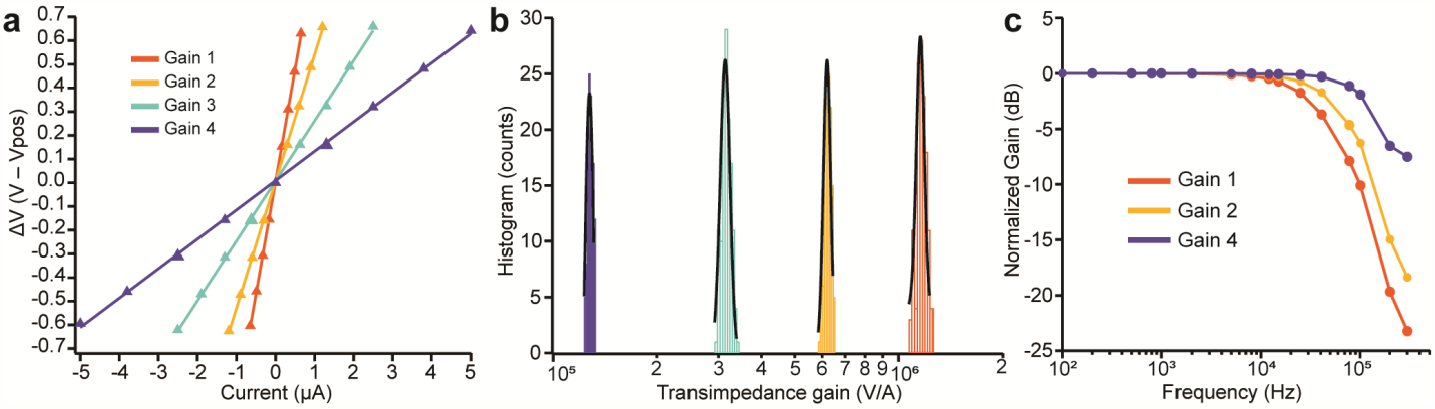
Measured transimpedance gain across the entire 128-amplifier array. (a) The gain calibration shows highly linear responses across the dynamic range shown with an R^2^ of 0.999 for all gain settings. (b) The 128-ch current sensing front-end exhibits a relatively consistent gain across the array with measured gains of 1.16 ± 0.04 MΩ, 618.93 ± 13.31 kΩ, 314.00 ± 9.63 kΩ, 127.27 ± 2.17 kΩ (mean ± SD) for Gain 1, Gain 2, Gain 3, and Gain 4 respectively. (c) Measured bandwidth of the TIA for Gain 1, Gain 2, and Gain 4 (measured cutoff frequencies of 40 kHz, 53 kHz, and 122 kHz, respectively).

### B. Bandwidth, noise, and crosstalk

To test the bandwidth of the amplifier, sinusoidal currents (100 Hz – 300 kHz) are injected into the amplifier while monitoring the response at the output (Fig. 4c). For the three gain settings tested, the cutoff frequencies are 40, 53, and 122 kHz for Gain 1, Gain 2, and Gain 4, respectively. Because the cutoff is determined by *R*_*f*_*C*_*f*_, the lowest gain setting (Gain 4) exhibits the widest bandwidth. Because in most cases, a constant bandwidth is desirable across gain settings, a second stage behaving as a low-pass filter can be placed to eliminate the variability. However, in this application, to minimize the footprint of each TIA, an additional low-pass filter is not used.

The noise charateristics are measured as functions of gain setting and biasing level. The output of the amplifier is sampled at 640 kS/s (Fig. 5). The noise level of the amplifier is 0.22 nA_RMS_, 0.42 nA_RMS_, and 1.99 nA_RMS_ for Gain 1, Gain 2, and Gain 4 respectively. For three different bias levels between 1. 1 μA up to 20.1 μA (Gain 1), no appreciable difference is observable in the noise of the amplifier (Fig. 5b).

**Fig. 5.**
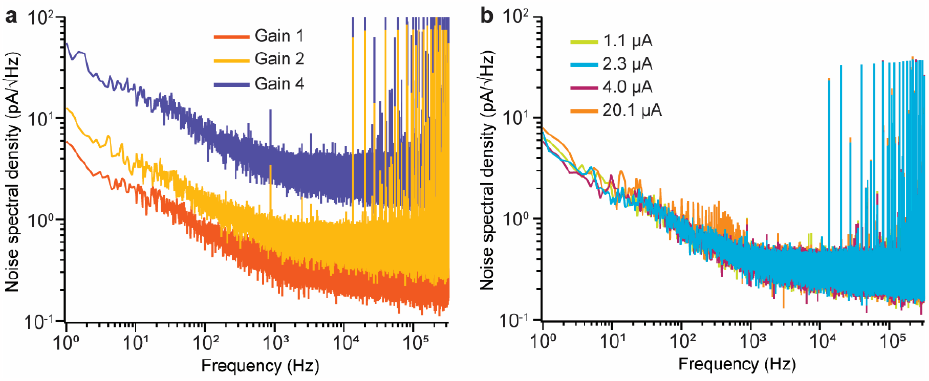
Noise characteristics in various TIA gain settings and biasing current. (a) The noise level of the amplifier is 0.22 nA_RMS_, 0.42 pA_RMS_, and 1.99 nA_RMS_ for Gain 1 (highest transimpedance gain), Gain 2, and Gain 4 (lowest transimpedance gain) respectively. (b) For bias levels between 1.1 – 20.1 μA at setting Gain 1, the noise density is consistent.

Crosstalk can often hinder the ability to provide independent high-density recordings. To verify the isolation between amplifiers in an array, it is important to examine crosstalk. Crosstalk is evaluated by introducing a large sinusoidal current (with a peak-to-peak amplitude of 0.8µA and a frequency of 625 Hz) that covers almost the entire input dynamic range into one amplifier and observing the spectral response of neighboring amplifiers (as shown in Fig. 6). The highest gain setting (Gain 1) is used for this experiment to test the worst-case scenario. In the time domain, the crosstalk between the physically-neighboring amplifiers is invisible (as shown in Fig. 6a). In the spectral responses, the crosstalk is measured at adjacent amplifiers and found to be -82.5 dB, -83.2 dB, and -89.7 dB (as depicted in Fig. 6b), showing nearly complete isolation between amplifiers.

**Fig. 6.**
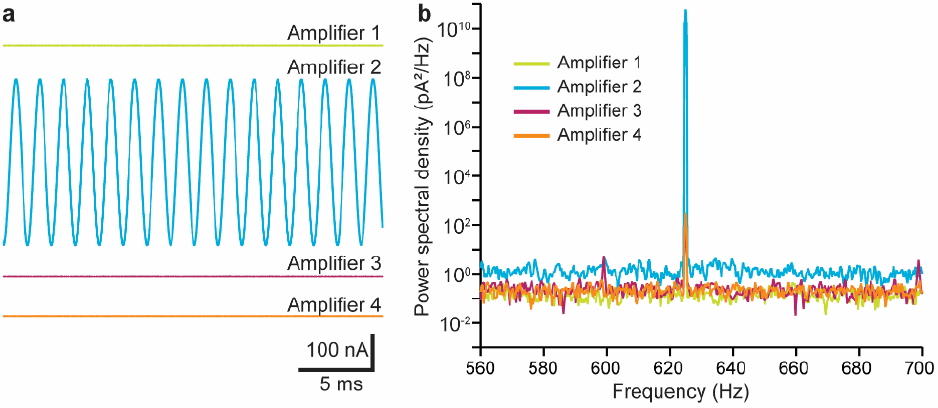
Crosstalk experiment with a sinusoidal current is injected into Amplifier 2. (a) In the time-domain, no crosstalk is observable. (b) The spectral analysis shows a small level of crosstalk in the adjacent amplifiers (−82.5 dB, -83.2 dB, and -89.7 dB).

## IV. Dopamine recordings using Carbon-fiber electrode

To validate the neurochemical-sensing front-end for dopamine measurement, one of the amplifier’s inputs is connected to a CFE. Using a flow cell setup, dopamine is introduced to the electrode while FSCV recordings are performed.

### A. Carbon-fiber electrode fabrication

The CFE fabrication process is based on a previously-reported method [9, 11]. In brief, a 7-µm carbon fiber is threaded in through a glass capillary using vacuum. Then, the glass capillary loaded with carbon fiber is pulled using a micropipette puller (P-87, Sutter Instument) which creates two glass pipettes with carbon fiber. The carbon fiber extending from the tip of the glass pipette is intially trimmed using a surgical scissor to a few milimeter in length. Subsequently, the glass pipette with carbon fiber is placed under a microscope and the carbon fiber is additionally trimmed using a surgical blade to ∼140 µm in length. The resulting carbon-fiber electrode has a sensing area of ∼3,100 µm^2^. The carbon fiber is connected to a 22 AWG wire using silicon epoxy in the opposite side of the glass capillary.

### B. Flow cell experiment

In order to introduce dopamine to the electrode with precision, a flow cell is typically used. For this experiment, a In-Vitro/FSCV microelectrode flow cell is used (NEC-FLOW-1, Pine Research). This flow cell is based on gravity feed. The fluid sample is loaded from below a flow channel (electrode channel) in which the carbon-fiber electrode is directly inserted. An Ag|AgCl reference electrode is used which is set to 1.55 V. This is to provide negative potential across the electrode-electrolyte interface without operating the CMOS chip at negative potentials. By default, phosphate-buffered saline (PBS) flows through the electrode channel with a flow rate of ∼1 mL/min. Dopamine solution can be added to this flow using a HPLC valve switch. This injection is governed by a syringe pump which controls the injection time, volume, and flow rate of dopamine solution (0.5 – 10 µM).

### C. Fast-scan cyclic voltammetry recordings of dopamine

All TIAs have a shared V_pos_ connection. The voltage ramp for FSCV is applied to V_pos_ node at each TIA. The scan rate is set to 400 V/s and the ramp range is between -0.4– 1.3 V (E vs. Ag|AgCl). Each FSCV period is set to 100 ms with a resting potential of -0.4 V.

During the recording session, the PBS buffer is used as a default solution with a continous flow rate of ∼ 1 mL/min. After 5 – 7 seconds of initial recording, DA solution is injected into the flow stream. For background subtraction, the initial 5-second FSCV recordings is averaged to create the background voltammogram.

For the 10-µM DA recording, the voltammogram taken after the DA injection shows distinct redox peaks compared to the background current (Fig. 7a). Using the background subtraction, redox peaks are visible at 400 mV and -100 mV (Fig. 7b), consistent with previous CFE-based DA FSCV recordings [9]. The presented CFE-based DA recordings validates the neurochemical-sensing front end for dopamine detection. The recorded cylic voltammograms can be plotted in a 2D image to visualize the oxidation and reduction of DA in time (Fig. 8a-b). The presence of dopamine can be seen between 7 – 20 seconds with strong oxidation and reduction currents (Fig. 8a). In a control experiment in which PBS is injected without DA, redox currents are not detected as expected (Fig. 8b). The oxidation peaks from individual cycles can be extracted to visualize the time-domain DA measurement. By extracting the oxidation currents at 400 mV, the transient response of DA show a rapid increase in the oxidation of DA at 7 seconds, which continues for ∼13 seconds before tapering down (Fig. 8c). No visible response is recorded in the control experiment (Fig. 8d), confirming that the current measured in the DA experiment is from the DA oxidation. Small ripplings in both 2D voltammograms are originating from 60 Hz noise which is likely due to the largely unshielded connection between the CFE and the amplifier chip. In practice, the amplifier chip will be used as a headstage amplifier chip that is in close proximity to the neural probe (<10 cm), and therefore, the 60 Hz noise will be substantially smaller.

**Fig. 7.**
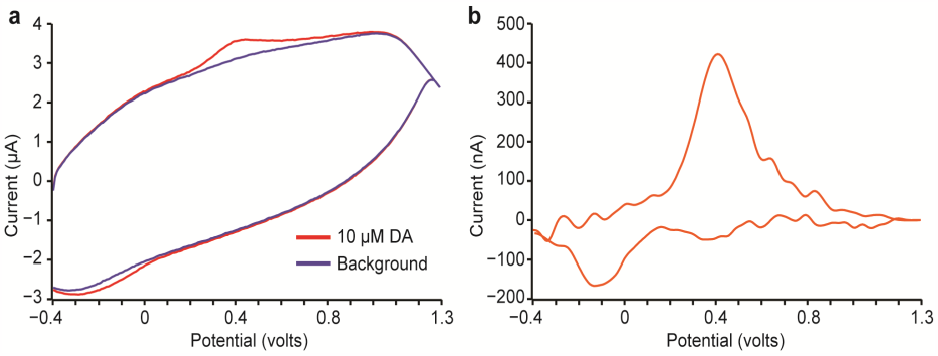
Fast-scan cyclic voltammogram using the neurochemical chip and an external CFE. (a) In the background measurement, the CFE measures a current signal in PBS without dopamine (DA). This measured current is mostly from the double-layer capacitance of the electrode-electrolyte interface. The measurement is repeated with 10-µM DA in PBS. Redox peaks are visible near 0.4 V and -0.2 V. (b) The 10-µM DA measurement is subtracted using the background measurement to show the redox reactions of DA.

**Fig. 8.**
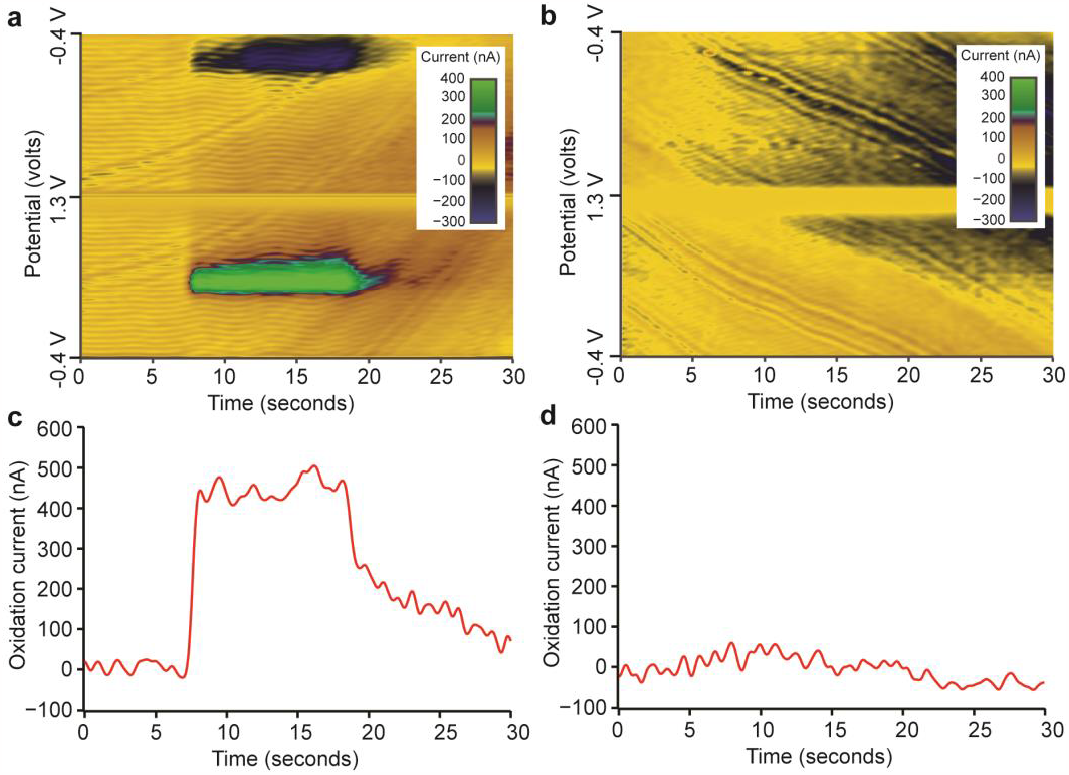
2D cyclic voltammogram using the neurochemical microchip and an external CFE. (a-b) The CFE is mounted on a flow cell where dopamine (DA) is introduced between 5 -7 seconds into the recording. For the control experiment, instead of DA, PBS is introduced. (c-d) the oxidation peaks of all the cycles are extracted for both DA measurement and control experiment.

In general, CFE should have a linear response to DA concentrations below 5 µM [25-27]. The flow cell experiment is repeated for various DA concentrations between 0.5 – 5 µM. The DA sensitivity of the CFE and the neurochemical-sensing front end is tested by plotting the oxidation currents for each DA concentrations (Fig. 9). Each experiment is repeated three times. The sensitivity is 51.9 nA/µM which is consistent with previous reports of DA detection using CFE [9].

**Fig. 9.**
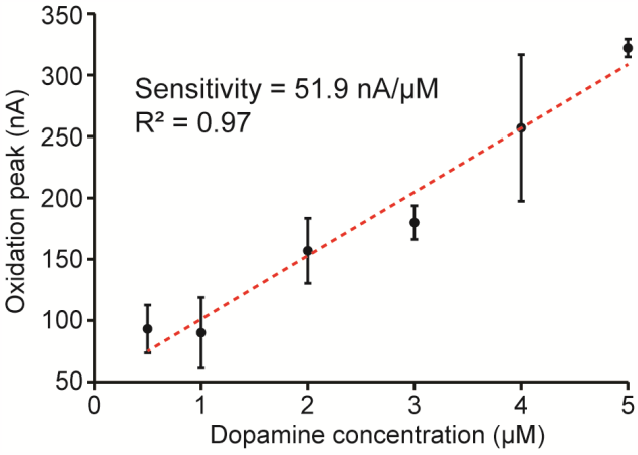
DA sensitivity of CFE and neurochemical-sensing front end. The error bar is in standard error of means.

### V. Conclusion and discussion

This paper introduces a high-density neurochemical microchip. In low-noise applications, an integrating capacitor-based TIA is a common approach. However, for FSCV applications, a wide dynamic range is required, which necessitates a substantial area investment for these integrating capacitors. This can be problematic when designing a large array of amplifiers. To address this, we investigated using n-well and poly resistors as feedback resistors to develop our scalable TIAs. The resistive TIAs we present have a small footprint of 220 µm × 80 µm and exhibit low-noise characteristics (0.22 nA_RMS_) that are suitable for *in vivo* neurochemical applications. The noise floor for the highest gain setting (BW = 40 kHz) is 1.1 pA/√Hz. We achieved a wide dynamic range of ± 5 μA, which can be expanded further by opting for a smaller resistance value, albeit at the expense of noise characteristics and resolution.

Compared to previously published works (Table I), our amplifier design achieves one of the lowest noise levels and smallest footprints. To quantify this works advantage, we defined a figure of merit (FoM) as the following:

**TABLE 1.**
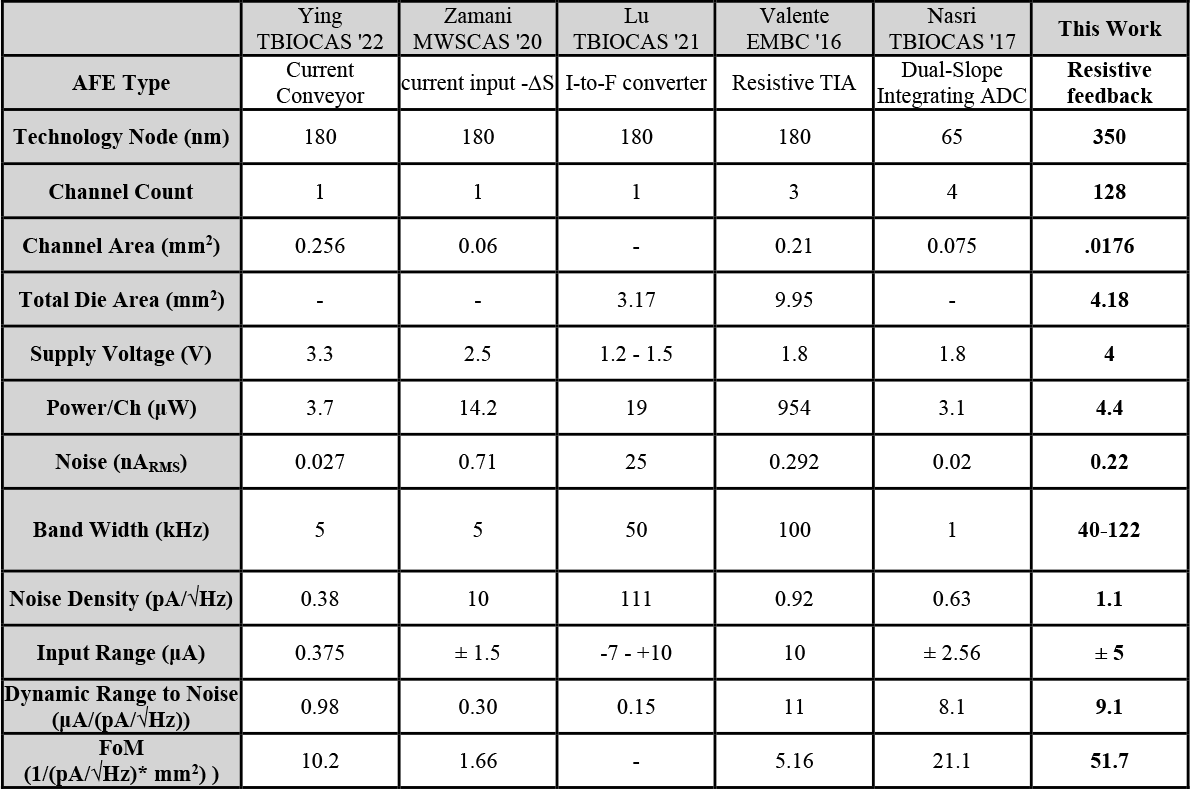
Comparison to State-of-the-Art current sensing front-ends

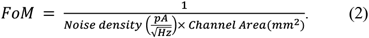

The FoM describes the area and noise efficiency for FSCV applications (higher the better). Because in large-scale neurochemical applications the area efficiency is critical, a large FoM indicates the quality of a scalable circuit topology for this application. Compared to previous research in Table I, this work achieves the highest FoM of 51.7. Our future goal is to assess the amplifier’s performance in FSCV applications by connecting it to a microelectrode array. This would enable high spatiotemporal *in vivo* imaging of electroactive neurotransmitters in the brain, a feat not yet accomplished.

One of the key challenges in developing high-spatial recordings of neurochemicals is the development of robust microelectrode arrays that have high level of specificity to faradaic current and stability without significant deterioration over time. While various materials have been tested for this purpose [28-32], carbon-based electrodes appear to be an appropriate candidate [32-34]. Our future work is aimed toward developing a microfabricated carbon-based electrode array in junction with this neurochemical chip.

*Appendix*

## Acknowledgment

Authors thank Advanced Microfabrication Facility (AMPAC) University of Central Florida staffs for their support. Also, authors thank University of Florida Research Service Centers with their support in microfabrication.

## Notes

### Competing Interest Statement

The authors have declared no competing interest.

